# *Anaplasma phagocytophilum* Subversion of Host Hepcidin-Ferroportin Iron Nutritional Immunity

**DOI:** 10.64898/2026.03.02.709072

**Authors:** Stephen L. Denton, Mingqun Lin, Elizabeta Nemeth, Yasuko Rikihisa

## Abstract

Obligatory intracellular bacterium *Anaplasma phagocytophilum* (*Aph*) causes an emerging infectious disease called human granulocytic anaplasmosis. Within host cells, *Aph* proliferates in a membrane-bound vacuole (*Aph*-vacuole), to which all required nutrients must be routed, including essential iron. Here, using a fluorescent labile iron-binding dye, we found that *Aph*-vacuoles are enriched with labile iron. Further, ferroportin (Fpn), the only known iron exporter in the plasma membrane of host cells, was increasingly localized to *Aph*-vacuole post-invasion.

Plasma membrane Fpn is tightly regulated by the hormone hepcidin, which binds to Fpn and triggers its ubiquitination and internalization. We found Fpn-GFP mutants deficient in ubiquitination (domain deletion or Y64H) had reduced or absent localization to *Aph-*vacuoles. Fpn-GFP (D131V) and Fpn-GFP (D039A) mutants that are deficient in iron binding and transport could still localize to *Aph-*vacuoles, but significantly reduced *Aph* growth and labile iron in *Aph*-vacuoles. *Aph* infection upregulated host cell expression of the hepcidin mRNA and protein. Furthermore, the gene encoding hepcidin is upregulated by inflammation, and we found *Aph* induced strong IL-6, IL-1β, and TNF-α mRNA expression in the host cells. Altogether, these results suggest that *Aph* induces production of proinflammatory cytokines for autocrine or paracrine hepcidin secretion to trigger the Fpn internalization, which can then be diverted to *Aph*-vacuoles in a ubiquitination-dependent manner as a mechanism of iron acquisition for the bacteria. This finding illuminates pathogen manipulation of cellular iron export mechanisms, and subversion of host hepcidin-Fpn iron nutritional immunity against pathogens.

**IMPORTANCE:** Iron is an essential element for both humans and microorganisms, serving as a cofactor in key metabolic processes. Obligatory intracellular bacterium *Anaplasma phagocytophilum* infects and proliferates within a membrane-bound vacuole (*Aph-*vacuoles) of neutrophils and endothelial cells, competitively acquiring intracellular iron from the host. Upon exploration of cytoplasmic labile free iron levels and distribution in the *A. phagocytophilum*-infected and uninfected host cells, we uncovered the unique ability of this bacterium to enrich intravacuolar iron. Ferroportin is the only cellular iron exporter of the host cells, which is regulated by the hepatic hormone hepcidin. Herein, we investigate the role of ferroportin and endogenous hepcidin locally produced by the host cells for iron enrichment in *Aph-*vacuoles. Ultimately, this study provides new insights into the novel mechanisms of microbial manipulation of host cells to acquire the essential micronutrient iron and overcoming host nutritional immunity, which may facilitate more effective treatment and prevention.

## INTRODUCTION

Rickettsioses including Anaplasmosis and Ehrlichiosis are among the most deadly vector-borne infectious diseases and greatly increasing in worldwide prevalence (1). Rickettsiae are obligatory intracellular bacteria that infect blood and endothelial cells. Human granulocytic anaplasmosis (HGA), caused by infection with *Anaplasma phagocytophilum* (*Aph*), is second in prevalence only to Lyme disease (2, 3), and human monocytic ehrlichiosis (HME), caused by *Ehrlichia chaffeensis* (*Ech*), is second in fatality only to Rocky Mountain spotted fever among tick-borne diseases in the US (4). *Aph* infects and replicates inside granulocytes (2, 5), and *Ech* infects and replicates inside monocytes-macrophages (6, 7). HGA and HME are severe flu-like febrile diseases accompanied by hematologic abnormalities and signs of hepatitis. There are no FDA-approved vaccines to protect against rickettsial diseases including HGA and HME. The only current therapy for HGA and HME is the broad-spectrum antibiotic doxycycline, which is generally effective with sufficiently early treatment, but no alternatives exist if cases are not treated early, or if microbial resistance develops.

Iron is an essential element for most living organisms, serving as a cofactor in key metabolic processes. As excess iron is toxic, however, systemic and intracellular iron levels are tightly regulated in animals by cellular iron import, storage, and export mechanisms (8, 9). Furthermore, sequestering iron from invading pathogens is an important arm of innate immunity, and is referred to as nutritional immunity (8, 10, 11). Cellular iron nutritional immunity is exemplified by the divalent cation transporter NRAMP1, that exports Fe^2+^ from late endosomes/lysosomes, driving resistance to facultative intracellular pathogens that occupy these niches (such as *Salmonella, Leishmania*, and *Mycobacterium*) by limiting pathogen growth (12). The known systemic iron nutritional immunity is hypoferremia mediated by peptide hormone hepcidin (Hepc, secreted mainly by liver), which not only deprives access to iron to blood-borne pathogens directly, but also shapes iron-dependent host cellular pathways that pathogens rely upon or defend against (such as glucose metabolism, oxidative stress, and immune regulation) (13).

For rickettsiae, iron acquisition is essential, but it is mostly unknown how these obligatory intracellular bacteria acquire iron from the host, and how they overcome host iron-sequestering nutritional immunity. While many other bacteria use siderophores, high-affinity iron-chelating compounds that competitively capture iron from the hosts, all rickettsiae members lack siderophores. *Ech* replicates in an early endosome-like compartment (vacuole) that does not fuse with lysosomes (14, 15). We previously discovered *Ech* acquires iron from 2 distinct mammalian iron-binding proteins: transferrin (Tf) and ferritin. Tf is the primary blood iron carrier protein. Host cells take up blood iron by endocytosis of iron-laden Tf that binds to the iron importer Tf receptor (TfR) on the cell surface (16). *Ech* subverts this pathway by driving upregulation of host cell TfR mRNA and subsequent recruitment of endocytosed Tf via the TfR to the *Ech*-containing vacuole ultimately gaining access to iron released from Tf (17–19). Consequently, interferon (IFN)-γ inhibits *Ech* infection by downregulating TfR (19). However, once infection is established *Ech* becomes resistant to IFN-γ (19), because *Ech* becomes capable of acquiring iron from ferritin (20). Ferritin stores iron in a stable, less reactive form until the cell needs it, so no pathogen was previously known to acquire iron specifically from ferritin. However, we discovered that *Ech* deploys via its T4SS (Type IV secretion system) the *Ehrlichia*-translocated factor effector Etf-3, which directly binds ferritin and induces ferritinophagy, degrading ferritin and conveying *Ech* access to liberated iron (20).

*Aph* is closely related to *Ech,* sharing 98% of protein coding-genes, and it requires iron (21). However, *Aph* replicates in the membrane-bound compartment with markers of early autophagosomes and ER-Golgi exit sites and lacks endosome-lysosome markers, the *Aph*-vacuole (14, 15, 22, 23). *Aph* does not use *Ech’s* iron-acquisition mechanisms, as unlike *Ech*, 1) *Aph* infection does not upregulate TfR mRNA (18); 2) *Aph-*containing vacuoles do not acquire Tf or TfR (14); and 3) *Aph* lacks Etf-3 orthologs that cause ferritinophagy. We, therefore, investigated *Aph* iron uptake mechanisms to create wider mechanistic understanding of iron-acquisition strategies used by rickettsial pathogens and enable developing new effective countermeasures.

## RESULTS

### Labile cellular iron is enriched in *Aph-*vacuoles, and ferritin is upregulated by *Aph* infection

Labile cellular iron (LCI) is the pool of Fe^2+^ iron in a cell that is not bound to storage molecules like ferritin. It is a crucial but transient intermediate in cellular iron metabolism, serving as a source of iron for metabolic processes. *Aph* infects only neutrophils, other granulocytes and endothelial cells *in vivo* (24). Human promyelocytic leukemia cell line HL-60 and monkey endothelial cell line RF/6A are relevant for *Aph* cellular and molecular biology investigations and have been widely used. As HL-60 cells are difficult to transfect, non-adherent, and round, RF/6A cells are used for transfection and unambiguous cellular localization, as they are flat and thinly spread adherent cells (25–27). To determine intracellular levels and distribution of the LCI pool in *Aph*-infected and uninfected cells, we used BioTracker 575 Red Fe^2+^ dye (28, 29) in live cell fluorescence microscopy, and ImageJ to quantitate fluorescence intensity in the area of a defined Region of Interest (ROI) (Supplemental Fig.S1). The result showed *Aph* has significant ability to enrich LCI pool in *Aph*-vacuoles in RF/6A cells (Fig. 1A-C). Consequently, total cellular LCI pool was significantly increased in *Aph*-infected cells than in uninfected cells (Fig. 1A & B). Cellular iron overload may increase intracellular ferritin because ferritin is the primary protein that stores and detoxifies excess iron within a cell (30). Thus, ferritin levels were determined by western blot analysis of ferritin light chain (FTL) normalized by human actin followed by densitometry. The result showed that in *Aph*-infected HL-60 cells, ferritin was significantly increased compared to uninfected cells (Fig. 1C & D). Ferritin synthesis is upregulated by signaling by proinflammatory cytokines, such as IL-6 (31), which is induced during *Aph* infection of human peripheral blood leukocytes (33) but not during *Ech* infection of human THP-1 monocytes (32). Altogether, This finding highlights opposite nature of *Aph*-infected cells vs. *Ech*-infected cells, wherein the latter ferritin is diminished due to ferritinophagy (20). Nevertheless, the enrichment of iron within *Aph-*vacuoles points to a distinct and novel mechanism of *Aph* infection on iron distribution and homeostasis within host cells.

**Figure 1.**
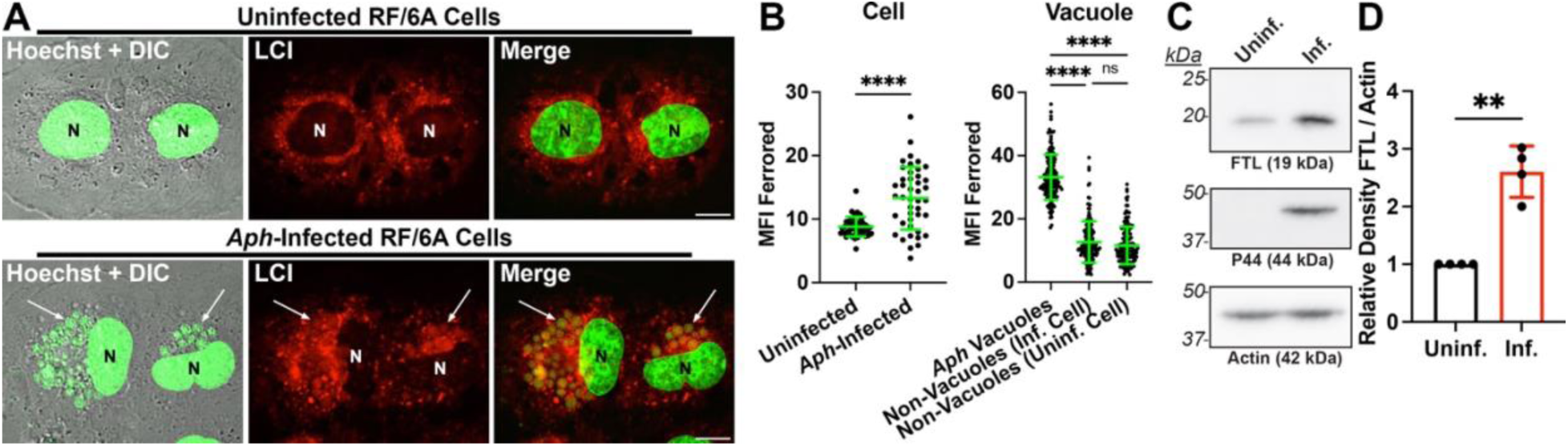
Labile cellular iron is enriched in *Aph-*vacuoles, and cellular ferritin is upregulated in *Aph*-infected cells. **(A)** LCI (Labile Cellular Iron, red) detected with BioTracker 575 Red Fe^2+^ dye in uninfected and *Aph*-infected RF/6A cells 1 dpi. Live cell image under a Leica Thunder microscope. DNA of nucleus (N) and *Aph* was stained with Hoechst dye (pseudocolored green). Small green globules indicated by white arrows are *Aph*-vacuoles. Merge, merge of the fluorescence images. DIC, differential interference contrast. Bar, 10 µm. (**B)** The scatter plots showing MFI (Mean Fluorescence Intensity) in arbitrary units in individual uninfected and *Aph*-infected cells, vacuoles and randomly paired non-vacuole areas with the horizontal bar representing the mean value (n = 43 cells/group, from which 153 vacuoles were assessed). Data are representative of two technical repeats and five total similar experiments. Student’s *t-*test (left graph) or One-Way ANOVA (right graph) where **** indicates *P* < 0.0001, ns, not significant. (**C)** WB analysis of uninfected (uninf.) and *Aph*-infected (inf. 2 dpi) HL-60 cells with anti-ferritin light chain (FTL), *Aph* P44, and actin antibodies. (**D)** Band density ratios of FTL relative to actin. The ratio of uninfected HL-60 cells was arbitrarily set to 1. Mean ± SD from four independent experiments. ** Significantly different by Student’s *t*-test (*P* < 0.01).

### Ferroportin (Fpn)-GFP and endogenous Fpn localize on *Aph*-containing vacuoles

Cellular iron levels are tightly regulated by iron import, storage, and export (8, 9). Fpn in the plasma membrane (PM) is the sole cellular Fe^2+^ exporter. Expression of Fpn on the PM leads to a decrease in LCI (33). Therefore, we analyzed intracellular distribution of Fpn-GFP in transfected RF/6A cells, and endogenous Fpn in HL-60 cells, with and without *Aph* infection. Both Fpn-GFP and immunofluorescence-labeled endogenous Fpn were distinctly localized on *Aph* vacuoles by tightly enveloping individual vacuoles (Fig. 2 A-D). Fpn-GFP were primarily localized on PM in uninfected RF/6A cells, whereas endogenous Fpn was homogeneously distributed in uninfected HL-60 cells (Fig. 2 E & F).

**Figure 2.**
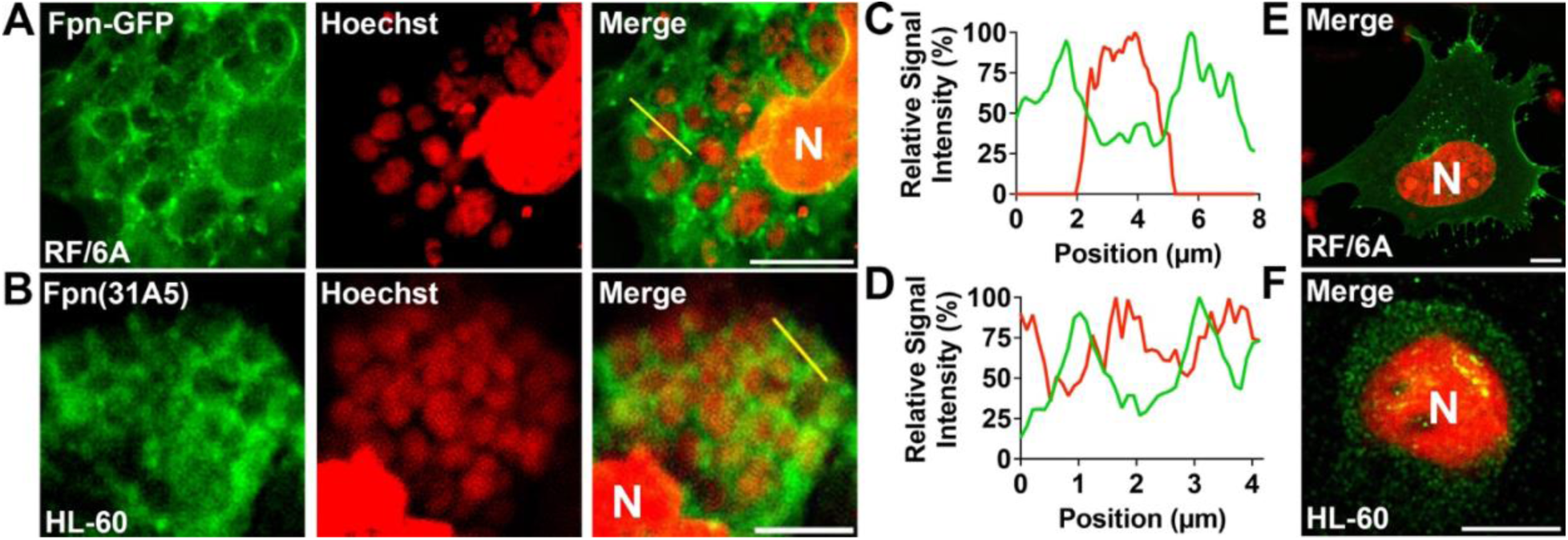
Fpn-GFP and endogenous Fpn localize to *Aph*-containing vacuoles. **(A)** Fpn-GFP (green)-transfected RF/6A cells were infected with host cell-free *Aph* at 1 dpt. At 3 dpt/2 dpi, cells were fixed and host and bacterial DNA were stained with Hoechst (pseudocolored red). Merge, merge of two fluorescence images. N, nucleus. White bar, 10 µm. (**B)** *Aph*-infected HL-60 cells were fixed at 2 dpi, permeabilized, and stained with mouse monoclonal anti-Fpn (31A5, green) and Hoechst. Bar, 5 µm. (**C, D)** Relative fluorescence signal intensity plots along the yellow lines depicted in A and B, respectively, of DNA (red line) with (**C**) transfected Fpn-GFP or (**D**) endogenous Fpn immunofluorescence labeled (green line), showing Fpn peaks encasing bacteria DNA peak in the vacuoles, but not overlapping. (**E, F)** Representative localization of Fpn-GFP at 2 dpt in uninfected RF/6A cells (**E**), or Fpn (31A5) immunofluorescence in uninfected HL-60 cells (**F**).

### Fpn-GFP colocalization with *Aph* vacuoles occurs subsequent to bacterial invasion and requires bacterial protein synthesis

Given Fpn-GFP and endogenous Fpn localization on *Aph* vacuoles occurred at exponential to stationary stages of bacterial proliferation (Fig. 2), next we examined early time course of Fpn localization. The results showed that at room temperature, *Aph* bound to RF/6A host cells, but did not internalize, and Fpn-GFP on PM did not colocalize with bound *Aph* (Fig. 3). However, at 37°C, *Aph* began to be endocytosed (Fig. 3), but Fpn-GFP did not start to localize on *Aph-*vacuoles until about 2 h post infection (hpi), and increasingly localized on the *Aph-*vacuole surface thereafter, showing a distinct ring-like pattern surrounding the vacuole (Fig. 3). This indicates that *Aph* binding alone is insufficient to induce PM Fpn endocytosis, and instead occurs upon *Aph* endocytosis, implicating an active process by intracellular bacteria. Thus, we assessed Fpn-GFP localization during late infection (2 dpi) with increasing periods of treatment with bacterial protein synthesis inhibitor, tetracycline. Starting at 2 h of treatment, *Aph* lost viability of continuous growth including drastic reduction of morulae (bacterial microcolonies) and disorganized colonies within the *Aph-*vacuole (Fig. 4A). Concurrently, the Fpn-GFP encasement of the *Aph-*vacuole became fragmented, diminished, or lost with tetracycline treatment (Fig. 4A) in treatment time-dependent manner (Fig. 4B). All together, these results indicate that both active intracellular bacteria and infection-induced host cell signaling must concur to induce PM Fpn endocytosis to eventually target *Aph* vacuoles.

**Figure 3.**
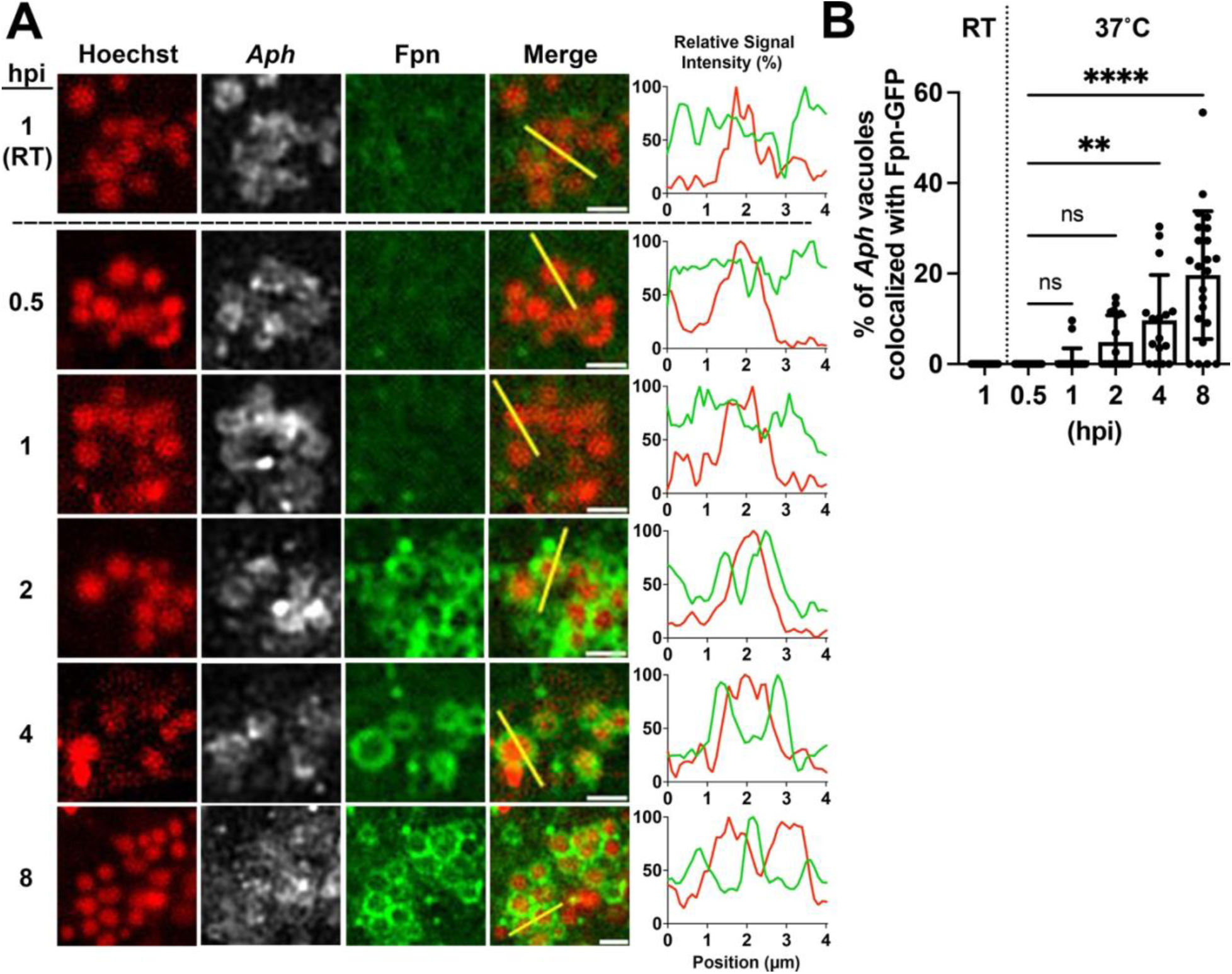
Fpn-GFP localization to *Aph* vacuoles occurs subsequent to bacterial entry. **(A)** Host cell-free *Aph* was added to Fpn-GFP-transfected RF/6A cells at 2 dpt at room temperature (RT) to allow only binding, or at 37°C to allow binding and internalization, and harvested at indicated hpi. *Aph* DNA was labeled with Hoechst (pseudocolored red) and *Aph* was labeled with anti-*Aph* (pseudocolored white). Merge, merge of Hoechst and Fpn-GFP images. White bar, 2 µm. Relative fluorescence signal intensity profiles between Fpn-GFP (green), and *Aph* DNA (red) along the yellow line in **A,** showing the Fpn peaks are encasing bacteria DNA peak within the vacuoles starting at 2 hpi. (**B)** Percentage of vacuoles encased with Fpn-GFP per cell. > 800 vacuoles or bacteria were counted across at least 20 cells at each time point. One-way ANOVA **** *P* < 0.0001, ** *P* < 0.01, ns, not significant.

**Figure 4.**
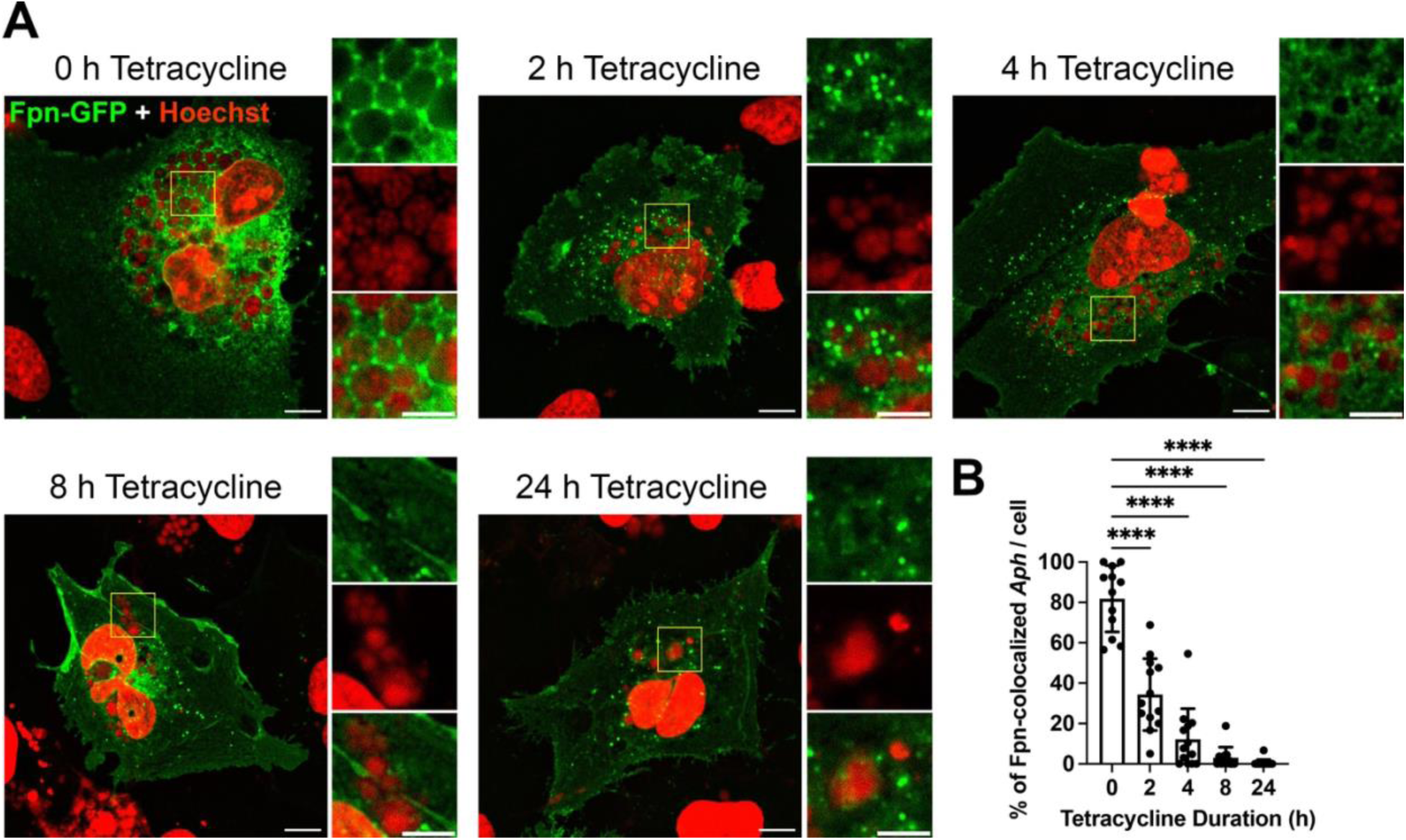
Fpn-GFP localization to *Aph-*vacuoles is dependent on *Aph* protein production. **(A)** RF6/A cells were transfected with Fpn-GFP for 2 d and infected with host cell-free *Aph* (MOI: 250) and treated with 5 µg/mL tetracycline for the indicated periods immediately proceeding harvest. Host and *Aph* DNA, labelled with Hoechst (pseudocolored red), and Fpn-GFP (green) fluorescence were imaged. Yellow boxes indicate inlaid area. White bar, 10 µm (whole image) or 5 µm (inlay) **(B)** % of *Aph* vacuoles with enveloping, non-fragmented Fpn-GFP rings scored in ImageJ. > 150 *Aph-*vacuoles were counted from 10 cells per treatment period. One-Way ANOVA to 0 h, **** *P* < 0.0001. Data representative of two independent experiments.

### Fpn motifs required for ubiquitination are necessary for Fpn-GFP localization to *Aph-*vacuoles

Fpn is a PM protein with 12 predicted transmembrane (TM) domains, and Fpn structure and function have been extensively studied in natural and experimental Fpn mutants, especially for the exogenous Hepc-dependent regulation of Fpn (34, 35). Hepc binding to Fpn causes rapid ubiquitination (Ub) of Fpn in cell lines overexpressing Fpn-GFP, and of endogenous Fpn in murine bone marrow-derived macrophages (36). Hepc-induced endocytosis of PM Fpn requires the 3rd (largest) cytoplasmic loop of Fpn, where multiple lysines (Lys) are localized and polyubiquitinated (36) (Supplemental Fig. S2). Thus, we determined if this host cell signaling pathway is usurped by *Aph* to internalize Fpn, without adding Hepc, using Fpn (Δ229-269)-GFP, Fpn (Δ225-247)-GFP, and Fpn (Δ247-269)-GFP(36). All these mutants can be transcribed, translated, and localize to the PM, but fail to show Hepc-induced internalization (36). Fpn (Δ229-269)-GFP cannot bind Hepc (36); Hepc-binding to other two mutants are unknown. Our results showed none of these partial deletion mutants of Fpn’s 3^rd^ cytoplasmic loop could effectively localize to *Aph* vacuoles (Fig. 5). To directly address if these Lys residues of Fpn in the 3^rd^ cytoplasmic loop are required for endocytosis to localize to *Aph-*containing vacuoles, we used Fpn(K8R)-GFP, in which 8 Lys between residues 229 and 269 in the 3rd cytoplasmic loop are substituted to arginine (34) (Fig. 5 and Supplemental Fig.S2). Fpn(K8R) binds Hepc normally, but cannot be endocytosed (34). In *Aph* infection, the Fpn(K8R) mutant also could not localize to *Aph-*vacuoles (Fig. 5). Lastly, Fpn(Y64H) is a natural point mutation in humans, that binds Hepc (34, 37), but in cells bearing it, Fpn is not ubiquitinated nor internalized (38), and ability to export iron is preserved (38, 39). We have created this mutant by site-directed mutagenesis of Fpn(WT)-GFP and found that Fpn(Y64H) also cannot localize to *Aph* vacuoles (Fig. 6). These results suggest Hepc-induced Fpn Ub pathway may be induced in *Aph*-infected cells without adding exogenous Hepc.

**Figure 5.**
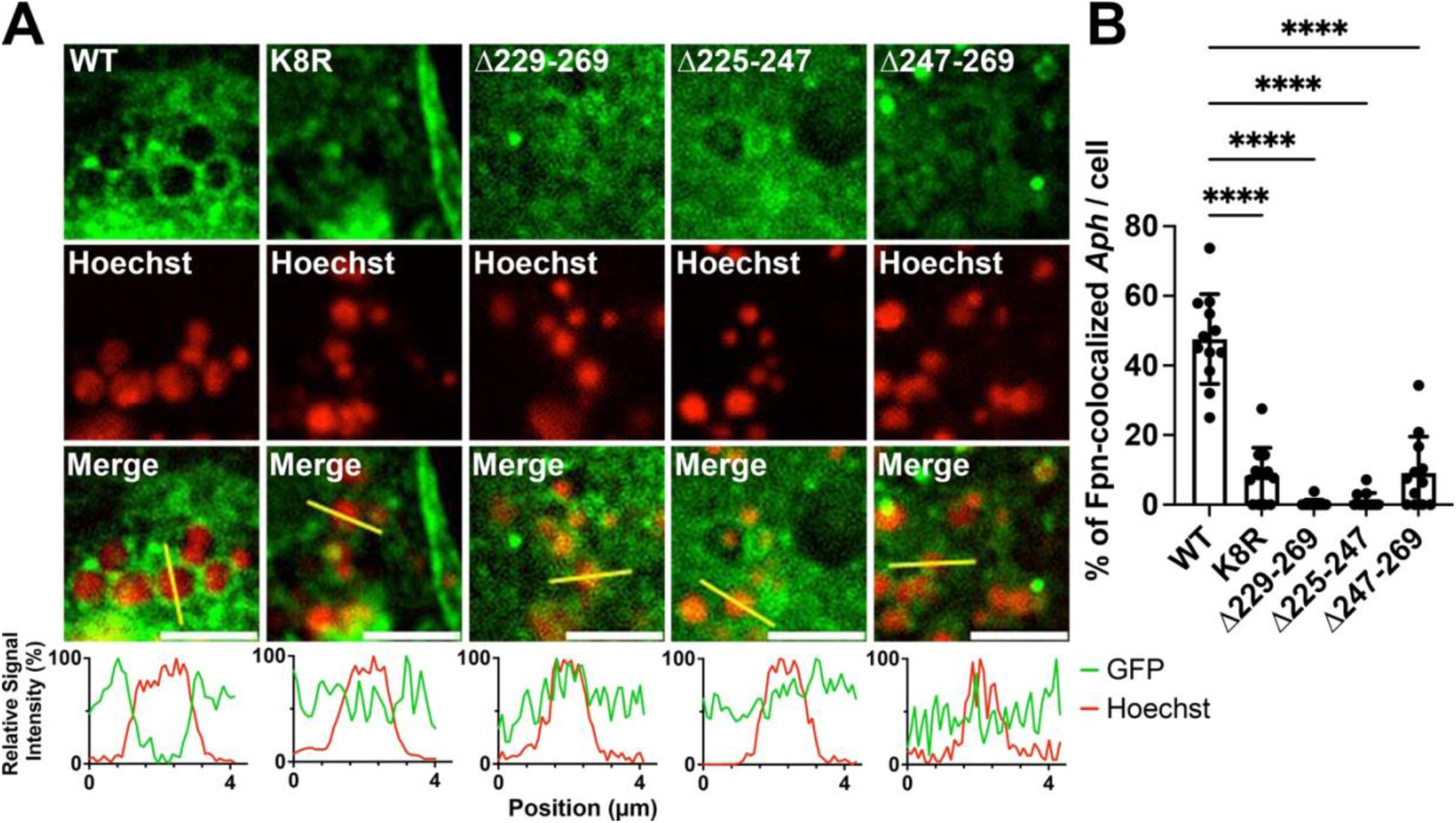
Fpn-GFP motifs for ubiquitination are required for Fpn-GFP localization to *Aph-*vacuoles. All lysine residues of Fpn-GFP (K8R) between positions 225-269 are substituted for arginine. This mutant along with Δ229-269, Δ225-247, and Δ247-269 are deficient in hepcidin-induced ubiquitination. **(A)** *Aph* was added to Fpn (WT or indicated mutant)-GFP (green)-transfected RF/6A cells at 2 dpt, and cells were harvested at 8 hpi. *Aph* DNAs were labeled with Hoechst (pseudocolored red). White bar, 5 µm. (**B)** Percentage of *Aph*-vacuoles encased with Fpn-GFP. > 200 vacuoles were scored in at least in 12 cells per condition. One-way ANOVA, **** *P* < 0.0001. Data are representative of three independent experiments.

**Figure 6.**
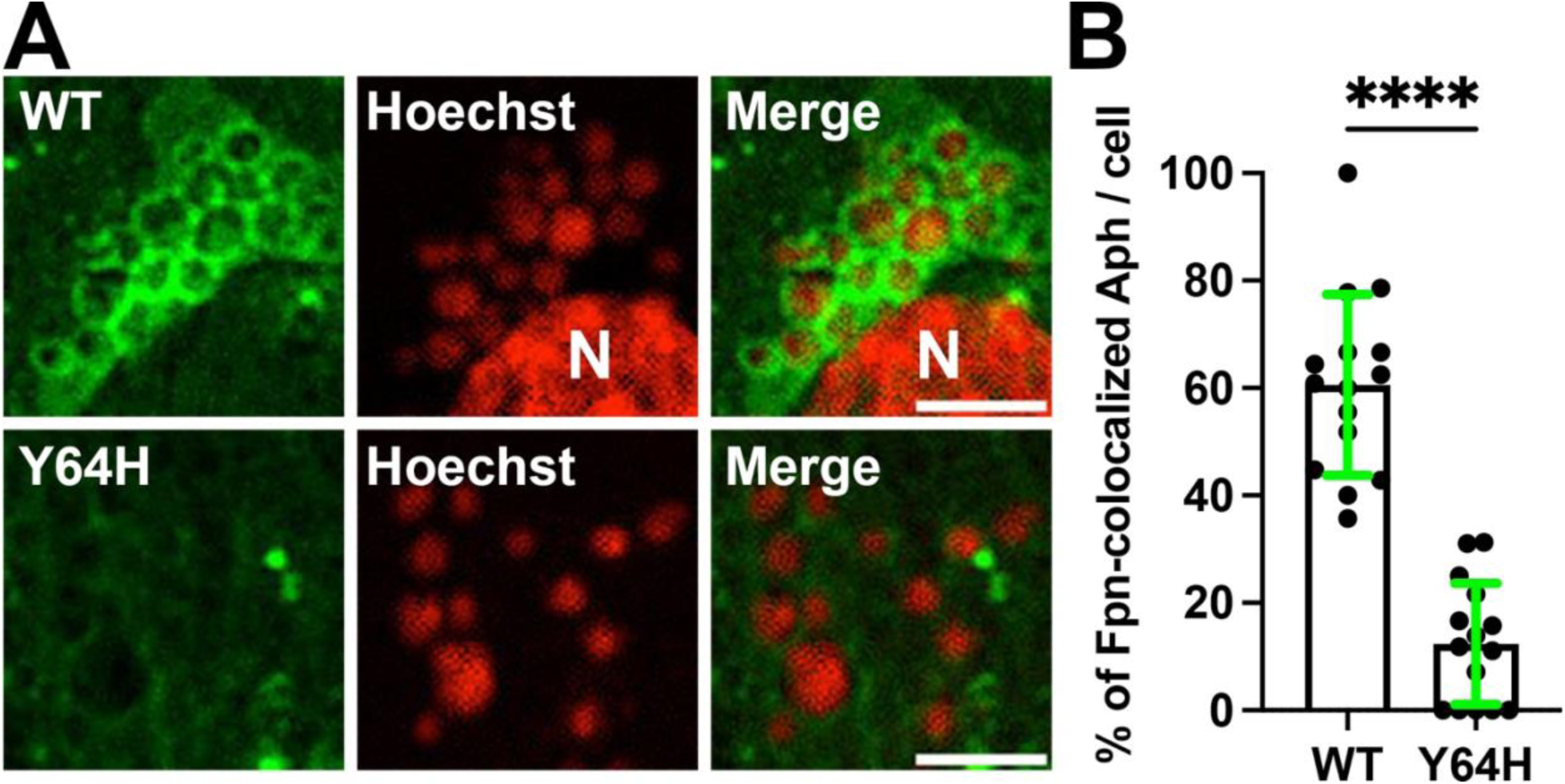
Fpn-GFP (Y64H), which binds hepcidin but cannot internalize, does not localize to *Aph-*vacuoles. **(A)** *Aph* was added to Fpn (WT or Y64H)-GFP (green)-transfected RF/6A cells at 2 dpt, and cells were harvested at 12 hpi. *Aph* and host DNAs were labeled with Hoechst (pseudocolored red). N, Nucleus. Bar, 5 µm. (**B)** Percentage of *Aph*-containing vacuoles with Fpn-GFP localization was scored by Image J. > 400 *Aph-*vacuoles were counted from 15 cells per condition. Student’s *t*-test, **** *P* < 0.0001. Data representative of two independent experiments.

### Iron binding/transport deficient mutants reduce LCI in *Aph* vacuoles and pathogen replication

The iron binding and transport sites of Fpn are in TM7, and conformationally coordinated at distal sites in the N-Lobe and C-Lobe (38, 40) (Supplemental Figs. S2 and S3). Fpn(D39A)-GFP cannot effectively bind ^55^Fe, and elevates the level of intracellular ^55^Fe *vs*. Fpn (WT)-GFP, due to impaired iron export (41). Fpn (D181V) and Fpn (N174I) are ^55^Fe binding and iron export-defective (41). Hepc-binding ability of these mutants are unknown. We, thus, tested if Fpn(D39A)-GFP or Fpn (D181V)-GFP can localize to *Aph* vacuoles, impair vacuolar LCI enrichment, and reduce *Aph* growth *vs*. Fpn (WT)-GFP by creating these Fpn mutants by site-directed mutagenesis. The results showed both Fpn(D39A)-GFP and Fpn (D181V)-GFP can localize to *Aph* vacuoles, like Fpn (WT)-GFP (Figs. 7B-D). Transfected expression of WT Fpn-GFP enhanced relative LCI levels in *Aph-*vacuoles/host cell cytoplasm compared to *Aph-*vacuoles of untransfected cells (UT), but not in Fpn (D039A)- or Fpn (D181V)-transfected cells (Figs. 7B-E). In turn, *Aph* growth within WT expressing cells was greater than in UT cells, a fitness benefit not afforded to *Aph* infecting D39A or D181V (Figs. 7F and G).

**Figure 7.**
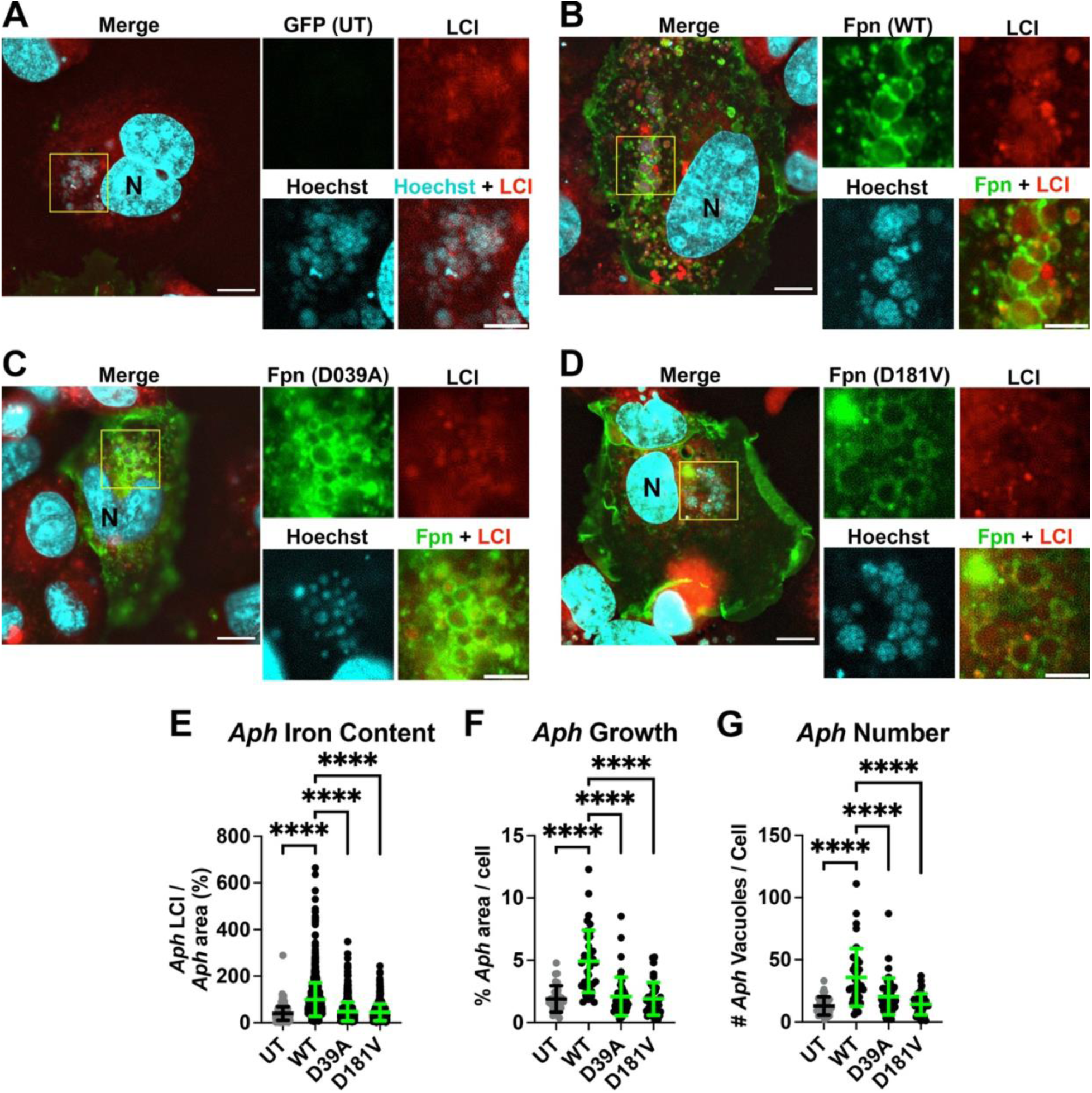
Iron binding/transport deficient mutants reduces LCI in the *Aph-*vacuoles and reduces infection. RF/6A cells were transfected with plasmids encoding WT, D039A, or D181V Fpn-GFP (colored green) and infected with host cell-free *Aph.* At 1.5 dpi and 3.5 dpt, cells were dyed with BioTracker 575 Red Fe^2+^ (Ferrored) to detect LCI (colored red) and Hoechst 33342 dye (pseudo-colored cyan) for detection of host and *Aph* DNAs. Live cells were imaged under a Leica DMi8 Thunder microscope at 37°C. Representative images are shown for cells: **A)** Untransfected (UT) **B)** Transfected with Fpn-GFP (WT), **C)** Transfected with Fpn-GFP (D039A), or **D)** Transfected with Fpn-GFP (D181V). Merge, merge of LCI, Hoechst, and indicated Fpn-GFP mutant. Inlays show individual fluorescent images of yellow-boxed areas. Bar, 10 µm, 5 µm within inlay. *Aph-*vacuole boundaries determined in ImageJ by globular, non-nuclear Hoechst fluorescence. **E)** *Aph* iron content measured by LCI MFI of vacuoles normalized by the *Aph* area % of the infected cell. N = 541-1324 vacuoles. **F)** % of *Aph* area / cell determined by area (µm^2^) of *Aph-*vacuoles / area (µm^2^) of occupied in GFP-expressing cell * 100. N = 30-45 cells. **G)** Total *Aph*-vacuole numbers were enumerated in GFP-expressing cells for each plasmid type. N = 30-45 cells. Statistics: One-Way ANOVA to WT control, ***p < 0.001 ****p < 0.0001. Data representative of three independent experiments.

### *Aph* infection induces production of autologous *HAMP* mRNA and Hepc protein levels

Infection with pathogens induces hepatocytes to produce and secrete Hepc into the blood(42). Hepc binds to Fpn in the PM of various cells, and induces Ub of Fpn, endocytosis, and thus lysosomal degradation(43), thereby reducing cellular Fe^2+^ export to the blood and preventing the appearance of non-transferrin-bound iron, the iron form which is critical for outgrowth of many invading extracellular pathogens (42, 44, 45). Hepc is synthesized as a prepropeptide with N-terminal endoplasmic reticulum targeting signal sequence, and upon cleavage by prohormone convertase furin produces the C-terminal 25 amino acid bioactive iron-regulatory hormone (mature peptide). Because Fpn was targeted to intracellular *Aph-*vacuoles even without adding exogenous hepcidin (Figs. 2-7), we examined if *Aph* infection induces endogenous Hepc production in HL-60 cells. The *HAMP* gene, encoding Hepc, is strongly upregulated in human hepatoma cells by proinflammatory cytokine IL-6{Nemeth, 2004 #8748}. The RT-qPCR result showed that *Aph* infection of HL-60 cells dramatically upregulated endogenous *HAMP* mRNA (Fig 8A). Western blot analysis using anti-human Hepc showed that *Aph* infection raises prohepcidin protein in HL60 cells (similar size as in (46, 47) (Fig. 8B-C). Concordantly, expression of proinflammatory cytokines TNF⍺, IL-6, and IL-1ß were significantly upregulated after infection, with the largest increase in IL-6 (Figure 8D). These results altogether strongly indicate that *Aph* subverts the Hepc/Fpn axis to directly provide iron and enhance fitness of this intracellular pathogen (Fig. 9).

**Figure 8.**
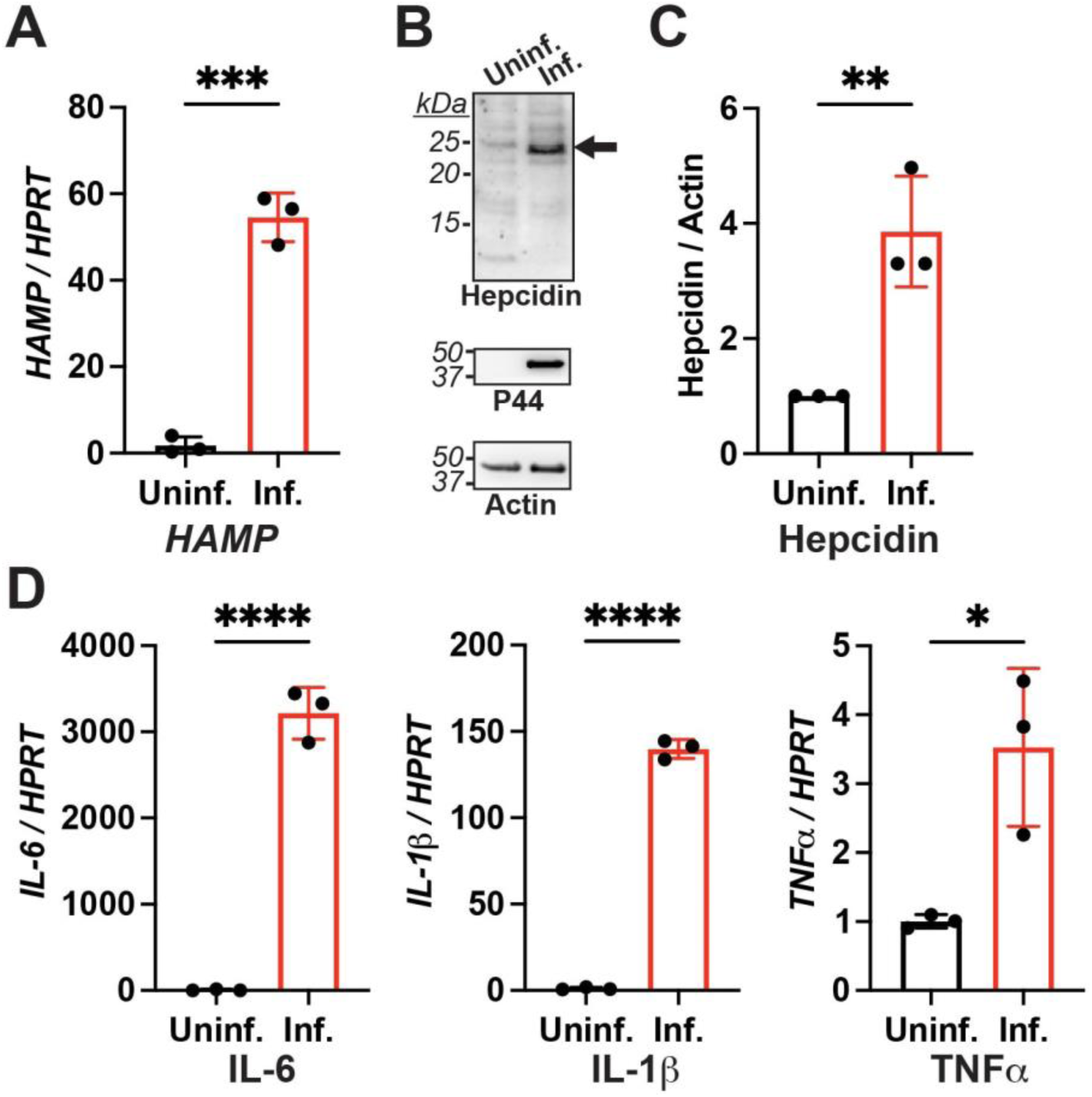
*Aph* infection induces production of Hepc by the host cells. **(A)** Relative *HAMP* mRNA levels normalized to human *HPRT* mRNA (ΔΔCt) in uninfected and *Aph*-infected HL-60 cells at 12 hpi. (**B)** Lysates from uninfected and *Aph*-infected HL-60 cells at 2 dpi were analyzed by western blotting using antibodies against human Hepc, human actin, and *Aph* P44. (**C**) The relative band density of Hepc was normalized against actin with the level of the uninfected group set as 1. **(D)** Relative mRNA abundance of human TNF⍺, IL_6, and IL-1ß from samples normalized to human HPRT as in (A) by the ΔΔCt method. Data indicate the mean ± SD from three independent experiments. Student’s *t*-test, ** *P* < 0.01; *** *P* < 0.001.

**Figure 9.**
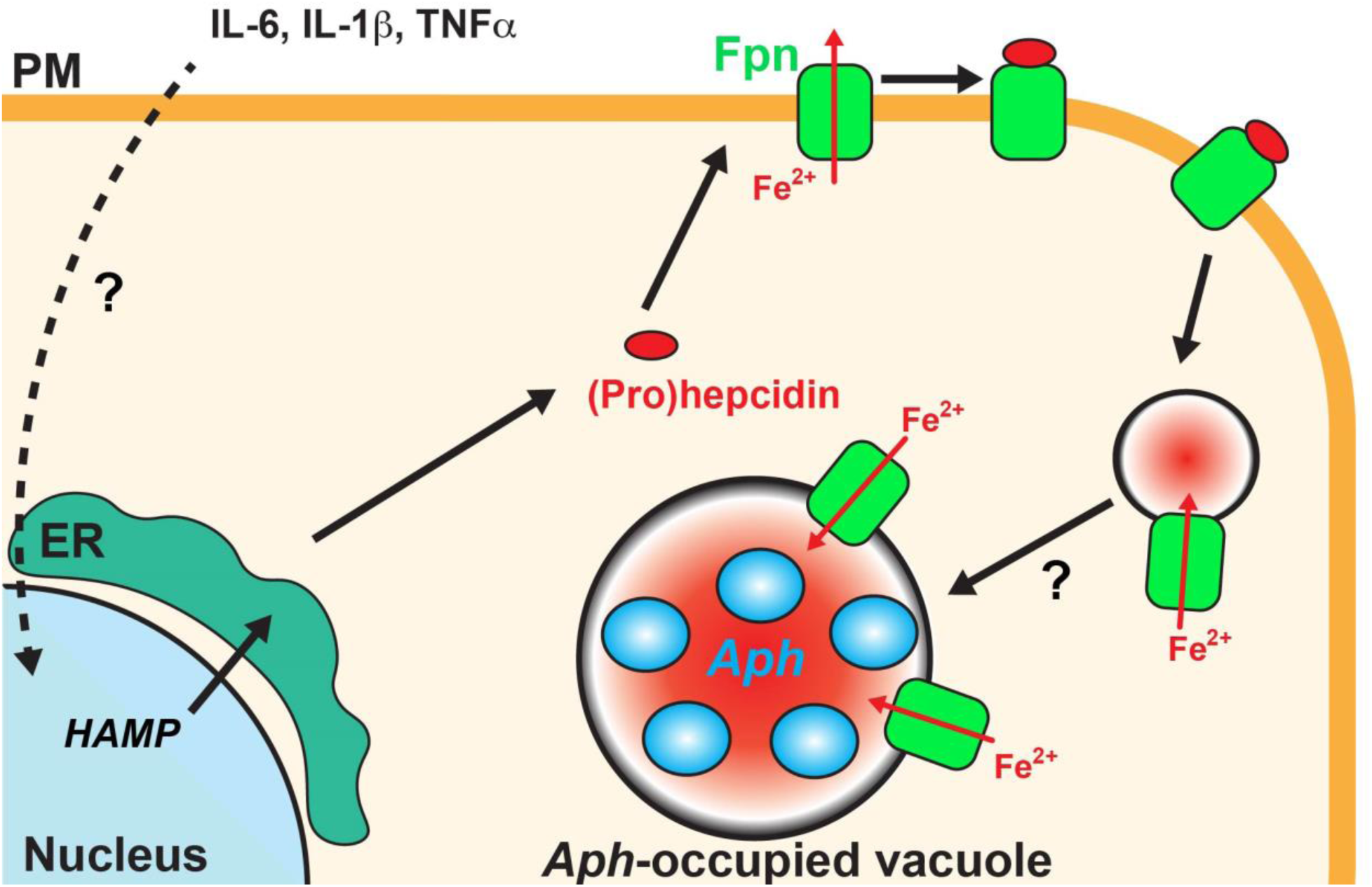
Hypothetical model: Infection-induced Hepc internalizes Fpn to *Aph* vacuole localization which facilitates iron uptake by *Aph*. PM Fpn exports iron from the cell. After *Aph* invasion, proinflammatory cytokines are upregulated concurrently with Hepc that induces endocytosis of Fpn, allowing Fpn endosomes to target *Aph-*vacuoles. Fpn localization on *Aph*-vesicles is vectorially positioned to transport Fe^2+^ from cytosolic sources into *Aph*-vacuole lumen, promoting bacterial growth. PM: plasma membrane, Fe^2+^: ferrous molecules, *Aph: Anaplasma phagocytophilum,* Fpn: ferroportin.

## DISCUSSION

Iron export by Fpn regulates systemic and intracellular iron homeostasis and is critical in nutritional immunity against pathogens (43). The present study revealed *Aph* recruits Fpn to its replicative vacuole to enable its proliferation. This contrasts with known prior mechanisms found in other microbes (10, 48). In some hereditary diseases, Fpn mutations cause organ iron overload (hemochromatosis) and increased risk of certain bacterial infections (48, 49). In macrophages infected with *Chlamydia psittaci, C. trachomatis*, or *Legionella pneumophila,* exposure to Hepc decreases Fpn, thus elevating intracellular iron, and enhancing their intracellular growth (10).

Fpn in the PM is co-internalized with *Staphylococcus aureus*, *Salmonella typhimurium* or IgG-coated beads when phagocytosed by macrophages, but then rapidly (< 15 min) removed from phagosomes and recycled back to the PM (50). Thus, two important iron-targeting host innate nutritional immunity mechanisms against intracellular infection are 1) maintaining or increasing PM Fpn levels to reduce total intracellular Fe^2+^, and 2) removing Fpn from pathogen-containing phagosomes. Contrasting with this established dogma, our data in two different cell lines revealed that Fpn was increasingly localized to *Aph* vacuoles. Iron transport-deficient Fpn reduced LCI in *Aph* vacuoles, consequently reducing *Aph* growth. As Fpn does not co-localize or co-endocytose with extracellular *Aph* and *Aph* protein production is required to maintain colocalization, a distinct signal for Fpn internalization and targeting to *Aph* vacuoles is generated by viable intravacuolar bacteria. Thus, our novel concept is that intracellular *Aph* subverts the Fpn-mediated iron restriction system, otherwise a potent mechanism of nutritional immunity, instead manipulating Fpn to traffic to its vacuoles so as to import Fe^2+^ into vacuoles for iron acquisition (Fig. 9). Molecular mechanisms for Fpn targeting to *Aph*-vacuoles remain to be elucidated.

During infection, inflammation drives Hepc upregulation in the liver to limit the availability of essential iron to invading pathogens, by blocking Fpn-mediated iron release from cells into the blood (51). In vitro stimulation of fresh human hepatocytes with a panel of cytokines showed strong induction of Hepc mRNA by IL-6, but not IL-1α or TNF-α (52), indicating that IL-6 is a primary proinflammatory mediator of hepcidin induction at a local level (53). Within 2 h after addition of *A. phagocytophilum*, IL-6, IL-1ß, and TNF-α mRNAs are induced in human peripheral blood leukocytes (PBLs) *in vitro* (54). Here, *Aph-*infection of HL-60 cells closely matched cytokine expression of infected human PBLs. Thus, Hepc upregulation in HL-60 cells by *Aph* infection may be a consequence of infection-induced inflammation and co-opted by *Aph* for Fpn internalization. Similarly, Ferritin acts as an acute-phase reactant, with levels rising sharply due to inflammation, infection, or cell damage, not only by iron overload (55). The present study revealed *Aph* is able to upregulate cellular ferritin light chain protein in HL-60 cells in the landscape of IL-6 upregulation, thus, infection-induced inflammation may have multifaceted roles in iron availability to *Aph*. A previous study, however, reported diminished ferritin complex level in HL-60 cells after *Aph* infection, despite an increase in ferritin mRNA (primarily heavy chain), as well as increases in ferritin complex and light chain mRNA in infected neutrophils (56). Reasons for these discrepancies are unknown. Interestingly, iron sensing by Iron-Responsive Protein-1 (IRP-1) activity in human monocytic leukemia cell line THP-1, is unchanged by *Aph* infection (18), suggesting impaired intracellular iron sensing and regulation of iron metabolism by IRP-1 may also be an important contributing factor in *Aph* infection.

Some bacterial pathogens induce Hepc by macrophages and neutrophils *in vitro* and *in vivo* in a toll-like receptor 4 (TLR4)-dependent manner (45). Undifferentiated HL-60 cells express low levels of TLR4 (59). Moreover, a defining trait of *Aph* is the absence of lipopolysaccharide (58), the ligand for TLR4. LPS-TLR4 axis is, therefore, unlikely to directly drive Hepc production in *Aph*-infected HL-60 cells. Recently, it was reported that TLR5 or TLR2:TLR6 heterodimer activation in murine primary hepatocytes induces *HAMP* mRNA(57). Again, as *Aph* also lacks TLR5 ligand, flagella, or TLR6 ligand, diacyl peptides (21), and expression of TLR2 in undifferentiated HL-60 cells is low (59), these pathways may not be involved in *HAMP* mRNA upregulation in *Aph* infection. Systemic *HAMP* gene transcriptional regulation involves signaling of iron levels and inflammation, both transcriptionally and post-transcriptionally. *HAMP* transcriptional activation requires the BMP/SMAD and HNF4α pathways and IL-6 activation of STAT3 (55, 58, 59) and stabilization of *HAMP* mRNA occurs through the HuR protein (60), but it is uncertain if these pathways are also involved locally in infected cells. Determining how intracellular *Aph* not only regulates Hepc production but also causes Fpn internalization and docking to *Aph* vacuoles remains an important question.

## MATERIALS AND METHODS

### *A. phagocytophilum, E. chaffensis*, and tissue cell culture

Cultivation of *A. phagocytophilum* (HZ) and *E. chaffensis* (Arkansas) in HL-60 cells (ATCC, Manassas, VA), preparation of host cell–free *A. phagocytophilum via* Dounce Homogenization, and infection were performed as described (61) (62). RF/6A monkey endothelial cells (ATCC) were cultured as described (27, 63). For experiments involving tetracycline, 5 µg/mL tetracycline (Sigma, St Lois, IL) was added to cultures at indicated timepoints.

### Plasmids

Plasmids encoding Fpn-GFP (WT) and Fpn-GFP mutants K8R, Δ229-269, Δ225-247, Δ247-269, and Δ547-562 were provided by Dr. Elizabeta Nemeth (36). Plasmids constructed in this study, primer sequences, and cloning strategies are described in Tables S1 and S2. Fpn-GFP mutants Y64D, D39A, and D181V were generated from Fpn-GFP (WT) using Q5 Site Directed Mutagenesis Kit (New England Biolabs, Ipswich, MA). Plasmids were purified with E.Z.N.A. Endo-free Plasmid DNA Mini Kit I (Omega Bio-tek, Norcross, GA), Endo-Free Maxi Kit (Qiagen, Hilden, Germany), or Nucleobond Midi EF Kit (Machery-Nagel, Düren, Germany).

### Antibodies

The mouse monoclonal antibody (mAb) 5C11 specific to *A. phagocytophilum* P44 was described previously (64). Horse antiserum immunoreactive with *A. phagocytophilum* (anti-*Aph)* from was preadsorbed with uninfected cell culture before use. The mouse anti-human Fpn (31A5) antibody was provided by Amgen, Inc. (Thousand Oaks, CA). The following antibodies were obtained from commercial sources – Santa Cruz Biotechnology (Dallas, TX): mouse anti-human B-actin (C4), mouse anti-GFP (B-2); Novus Biologicals (Centennial, CO): rabbit anti-human Fpn (21502); Alpha Diagnostics International (San Antonio, TX): rabbit anti-human Hepc (HEPC13-S). Secondary antibodies including Alexa Fluor 488 (AF488)-conjugated goat anti–mouse IgG was obtained from Thermo Fisher, Cy3-conjugated goat anti-horse Ig was obtained from Jackson Immunoresearch, and peroxidase-labeled goat anti–mouse or rabbit secondary antibodies were obtained from Seracare (Milford, MA).

### RT-qPCR

At 2 dpi, cells were washed with phosphate-buffered saline (PBS; 8 mM Na_2_HPO_4_, 1.47 mM KH_2_PO_4_, 2.67 mM KCl, 137.9 mM NaCl, pH 7.4), and RNAs were extracted with the RNeasy Mini kit (Qiagen, Germantown, MD). cDNA was synthesized using the Maxima H minus First Strand cDNA Synthesis kit with Oligo-dT primers (Thermo Fisher Scientific), and bacterial and host gene expressions were determined by RT-qPCR with specific primers for *Aph* 16S rRNA (65), human genes *HPRT* (this study), *Fpn* (66), *HAMP* (67), *TNF⍺* (68), *IL-1ß* (69) and IL-6 (70) (Table S2), using Maxima SYBR Green/ROX Master Mix (Thermo Fisher Scientific) in a AriaMx Real-time PCR system (Agilent, Santa Clara, CA) according to the manufacturers’ instructions.

### Western blot analysis

Uninfected or *Aph*-infected cells at 2 dpi were washed with phosphate-buffered saline (PBS; 8 mM Na_2_HPO_4_, 1.47 mM KH_2_PO_4_, 2.67 mM KCl, 137.9 mM NaCl, pH 7.4) and soluble protein was extracted using commercial mammalian protein extraction reagent (M-PER, ThermoFisher Scientifc #78501). Following high speed centrifugation, soluble protein was reduced with 5X loading buffer (30% Glycerol, 2.3 M 2-mercaptoethanol, 4% w/v SDS, 0.03% bromophenol blue, 250 mM Tris-HCl pH 6.8) and denatured by boiling for 5 minutes. Proteins were separated by 10% (Fpn, Actin, and P44), 12% (FTL), or 15% (Hepc) SDS-PAGE gel electrophoresis followed by semi-dry transfer to 0.45 µm nitrocellulose (Fpn, Actin, P44, and Hepcidin) or methanol-activated 0.2 µm PVDF (FTL). Protein loads were normalized by BCA Assay or equivalent and verified by Ponceau S staining of the membrane and anti-actin immunoblots. Membranes were blocked and blotted with 5% skim milk and 0.05% Tween-20 in tris-buffered saline (TBS, 15 mM NaCl and 5 mM Tris, pH 7.4) and developed with ECL Detection Reagent (ThermoFisher #1859701). Chemiluminescence was photographed with a GE Amersham Imager 680 and band densities recorded in NIH ImageJ.

### Transfection, immunofluorescence (IF) labeling, and cellular localization analysis

RF/6A cells adhered to coverslips were transfected using FuGENE HD reagent (Promega, Madison, WI) with plasmids at 3:1 ratio (3 μl FuGENE HD : 1 μg plasmids) in a 12-well plate according to the manufacturer’s instructions. Cells were then infected with host cell free *Aph* at MOI 500 at timepoints indicated. In some experiments, cells were infected by co-culture with highly infected HL-60 cells for one day prior to transfection. For IF labeling, cells were fixed and permeabilized with ice cold 80% methanol/20% acetone, and labeled with primary then fluorescently-conjugated secondary antibodies diluted in 0.05% bovine serum albumin (BSA, Sigma-Adrich). In some experiments, cells were fixed with 4% paraformaldehyde (PFA, Sigma) and labeled with primary and then fluorescence-conjugated secondary antibodies diluted in PBS supplemented with 0.1% gelatin (Bio-Rad, Hercules, CA), 0.05% bovine serum albumin (Sigma), and 0.1% saponin (Sigma). Uninfected HL-60 cells or *Aph-*infected HL-60 cells were cytocentrifuged onto a microscope slide and fixed with ice cold 80% methanol / 20% acetone. Labelling occurred in a humidification chamber with primary and then secondary antibodies diluted in 0.05% BSA in PBS. In all experiments, the host cell nuclei and *A. phagocytophilum* DNA were labeled with Hoechst 33342 (Thermo Fisher Scientific).

### LCI analysis

For live cell LCI analysis, RF/6A cells were adhered onto glass-bottom tissue culture dishes (CellVis, Mountain View, CA) and infected with host cell-free *Aph* for two days. LCI was detected after incubation with 5 µM BioTracker 575 Red Fe^2+^ dye (Sigma-Aldrich) for 1 h at 37°C with 5% CO_2_ using an Oko Labs incubation chamber microscope mount. Nuclei and *Aph* were labeled with Hoechst, then live cells observed under Leica Thunder Imager at 37°C (20). For LCI analysis of *Aph* occupying Fpn(WT)-GFP, Fpn(D39A)-GFP, and Fpn(D181V)-GFP-transfected RF/6A cells, cells were infected with host cell-free *Aph* (MOI: 500) at 1 dpt. At 1.5 dpi (2.5 dpt), cells were incubated with 5 μM BioTracker 575 Red Fe^2+^ dye at 37°C for 1 h.

### Image analysis

Fluorescence images with overlaid differential interference contrast (DIC) images were acquired and analyzed with a Leica Thunder imaging system with computational clearing to remove any out-of-focus background (Leica Microsystems, Deerfield, IL). NIH ImageJ software was used to measure line scan relative signal intensities or area mean fluorescent intensities. A macro (described in Supplemental Fig. S1) was written to randomly locate and calculate the area of a defined ROI dimensions of each *Aph*-vacuole to measure mean fluorescence intensity (MFI) of “non-vacuoles” from infected or uninfected cells within the same field of view.

### Statistical analysis

Statistical analysis was performed with Student’s unpaired *t* test or a one-way analysis of variance (ANOVA). *P* < 0.05 was considered statistically significant. All statistical analyses were performed using Prism 10 (GraphPad, La Jolla, CA).

## Acknowledgments

This study was funded in part by a R01AI151065 grant to YR from the National Institutes of Health.

## SI Appendix

Supporting information for this article, including Supplementary Methods, Figures, and Tables can be accessed on the publisher’s website.

