## Supplementary Information for "*Anaplasma phagocytophilum* Subversion of Host Hepcidin-Ferroportin Iron Nutritional Immunity"

**Running Title:** Ferroportin manipulation by *Anaplasma*

**KEYWORDS:** *Anaplasma phagocytophilum*, ferroportin, iron transport, hepcidin

**Figure S1. LIP measurements.**

For experiments of fluorescent measurement of the labile iron pool (LIP) in live cells, images were collected and BioTracker 575 Red $Fe^{2+}$ mean fluorescent intensity (MFI) of regions of interest (ROI) were measured with ImageJ (NIH). A) Example ROI demarcation in a merged Hoechst, BioTracker 575 Red $Fe^{2+}$, and DIC image. For cell measurements (Fig. 1B), cell boundaries were determined in the DIC channel and classified as uninfected (white boundary) or *Aph*-infected by the presence of at least one non-nuclear vacuole containing bacterial DNA (yellow boundary). For vacuole measurements (Fig. 1B), within infected cells the boundaries of *Aph*-vacuoles were determined by round bacterial DNA clusters in the Hoechst channel (green). For each *Aph*-vacuole, the ROI Random Redistributor macro (shown in panel B), was applied to move the exact dimensions of the ROI to random locations in the image. If the random redistribution was contained within an infected cell boundary, it was classified as an infected cell non-vacuole (blue), and if it fell into an uninfected cell boundary, it was classified as an uninfected cell non-vacuole (magenta). All MFI measurements were taken in the BioTracker 575 Red $Fe^{2+}$ channel.

**Figure S2. 2-Dimensional representation of Fpn structure interpreted from Protein Database**

Illustration adapted from concept in (36) and based on Protein Database (PBD) crystal structures 5AYN, 5AYO, 8C03, 8BZY, 6W4S, 6WBV, and 6W4V (40, 71, 72). Amino acids with dotted lines have not been resolved in reported crystal structures, whereas solid lines have been resolved and flattened circles represent alpha helical structure. Yellow designates residues that are involved in iron binding or export function (41, 73, 74), and green designates residues involved in Hepc binding (37) (34, 36, 38, 39). Pink residues indicate lysines that have been mutated in the K8R multiple mutation of Fpn-GFP and intensely colored if ubiquitinated by hepcidin (34). Circled regions have been deleted in Fpn-GFP mutants and are color matched to the residue number(36). The epitope of antibody 31A5 is shown.

**Figure S3. Molecular mechanism of ferroportin-mediated iron export and hepcidin occlusion**

Illustration interpreted from structural and functional analysis reports (34, 36-41, 71-74). Fpn is made up by twelve alpha-helical transmembrane domains (TM1-12) divided into two distinct lobes according to their amino and carboxyl termini, named the N-Lobe (TM1-6, orange and yellow, colored to differentiate TM domains) and the C-Lobe (TM7-12, blue and green). Further features include a perpendicularly oriented alpha helical "hinge" anterior to TM7, a disordered intracellular domain between the hinge and TM6, and a disordered extracellular domain between TM9 and TM10. Ferroportin is crystalized in two distinct conformations according to the exposed orientation of the iron/hepcidin-binding pocket: the inward-facing state and the outward facing state (40, 71, 72). In both conformational states, the transmembrane domains are divided into two lobes, according to their amino and carboxyl termini orientation: the N-Lobe made up of interweaving TM1-6 (colored orange and yellow) and the C-Lobe made up of interweaving TM7-12 (colored blue and green). Conformational states differ dramatically in positioning of the two alpha helices comprising TM7, denoted as TM7a and TM7b. Accordingly, TM7 is heavily composed of residues critical for iron-binding, export, and hepcidin binding (38, 39). Intracellular ferrous iron has access to the inward-facing state of ferroportin, which triggers a ligand-induced conformational change to the outward-facing state, opening the binding pocket to the extracellular environment for iron export and hepcidin binding involving interactions of other transmembrane domains TM1 (including aa D39 and Y64), TM5 (including aa D181), and TM10

(38, 40). Hepcidin binding ferroportin in the outward facing state both occludes the pocket, preventing iron export, as well as initiating the ubiquitination-internalization-degradation cascade (34, 36, 43).

A

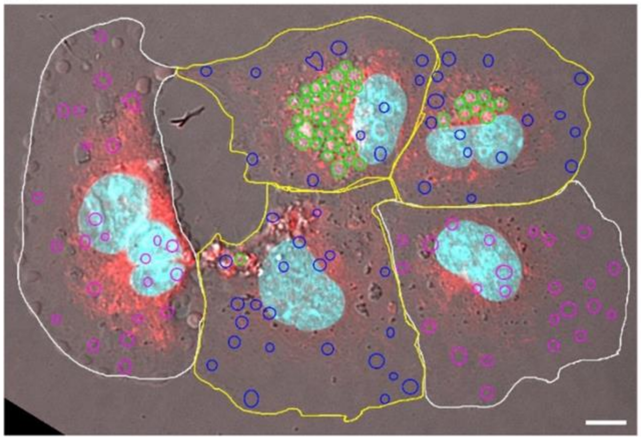

B

```
macro "ROI Random Redistributor [g]" {
a = getWidth();
b = getHeight();
x = (random) * a;
y = (random) * b;
Roi.move(x, y);
}
```

**Figure S1. LIP measurements.**

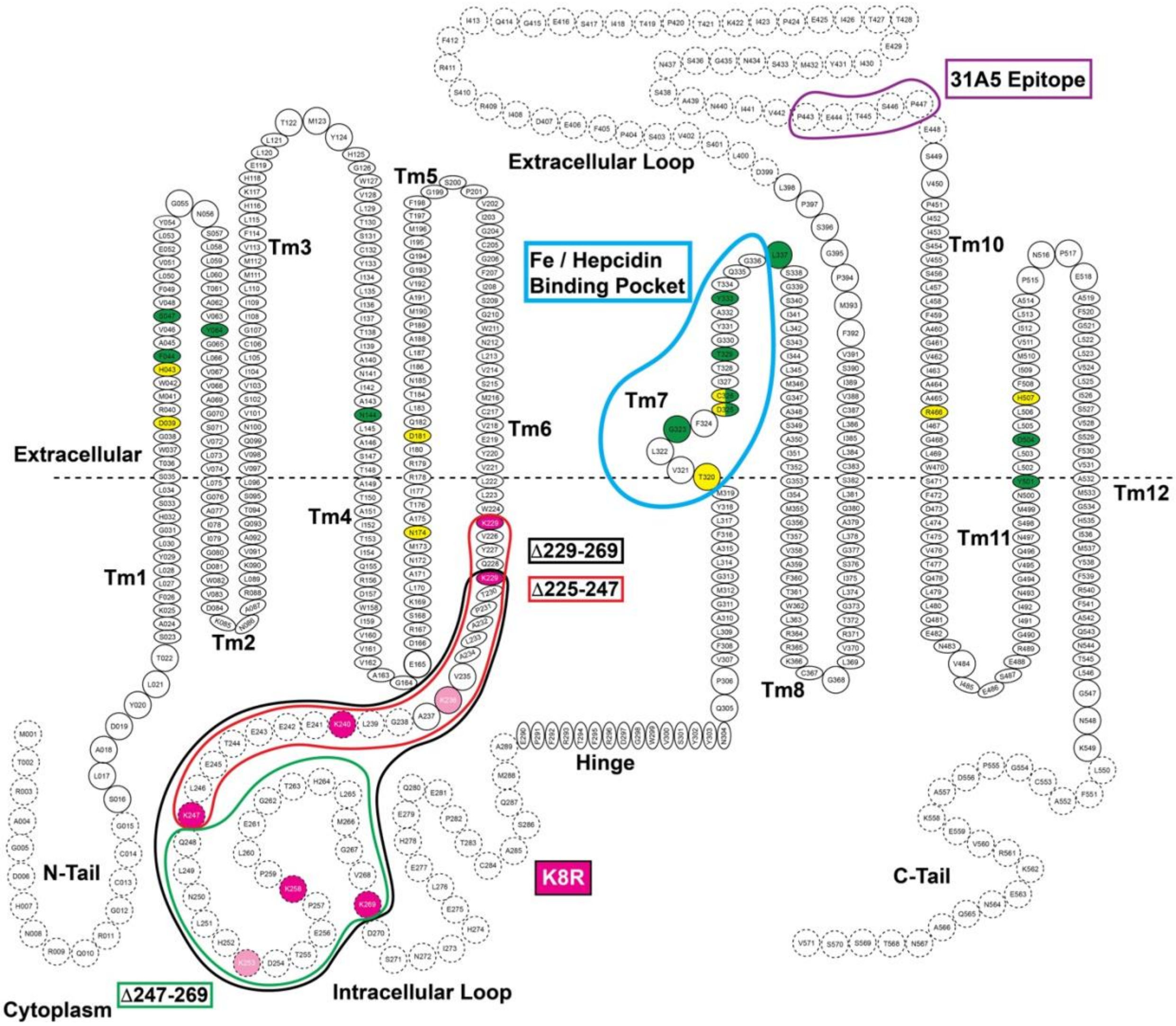


**Figure S2. 2-Dimensional representation of Fpn structure interpreted from Protein Database**

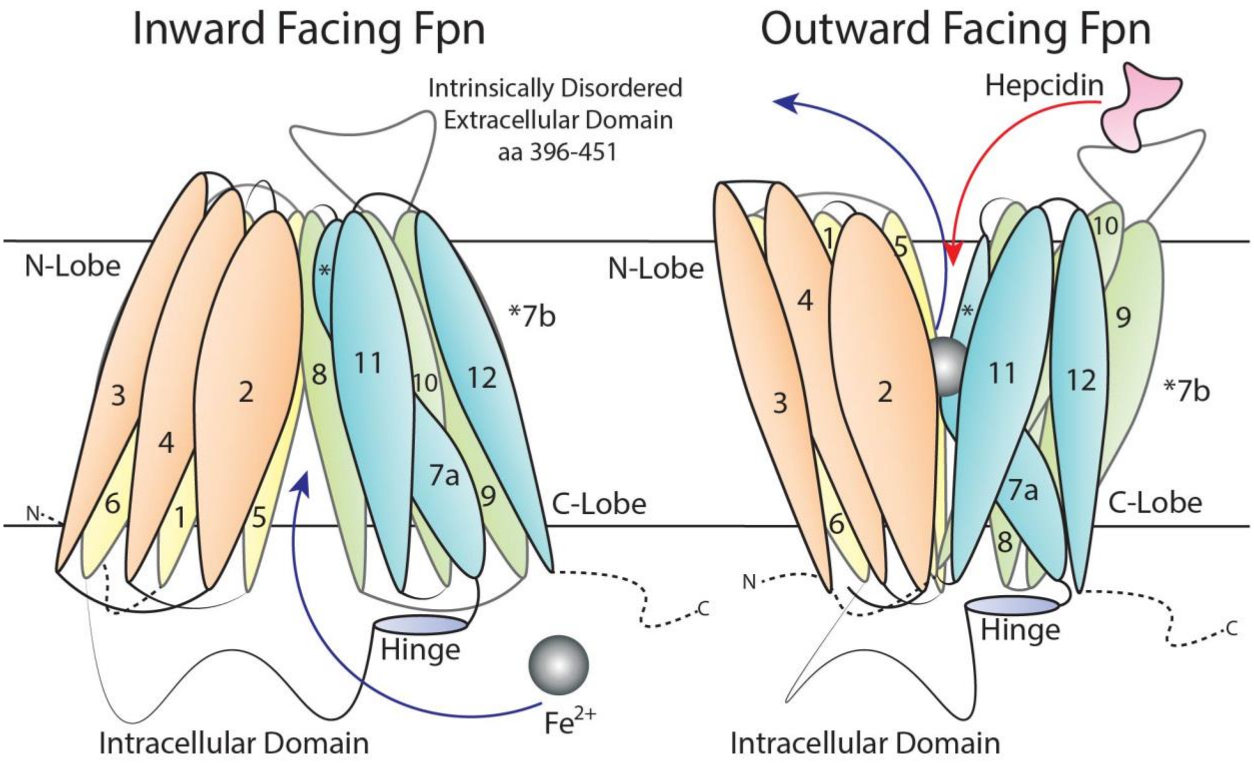


**Figure S3. Molecular mechanism of ferroportin-mediated iron export and hepcidin occlusion**

**Table S1.**

| Plasmid Name | Description | Reference |
|---|---|---|
| Fpn-GFP (WT) | Functional Human Fpn with C-Terminal EGFP Tag. | 38 |
| Fpn-GFP (Δ229-269) | C-terminal EGFP-tagged Fpn lacking residues 229-269 that make up the intracellular loop. Kanamycin selection. | 36 |
| Fpn-GFP (Δ225-247) | C-Terminal EGFP-tagged Fpn lacking residues 225-249 that make up the intracellular loop | 36 |
| Fpn-GFP (Δ247-269) | C-Terminal EGFP-tagged Fpn lacking residues 249-269 that make up the intracellular loop | 36 |
| Fpn-GFP (K8R) | C-Terminal EGFP-tagged Fpn where the 8 lysine residues of the ntracellular loop have been mutated to arginine. | 34 |
| Fpn-GFP (Y64H) | C-Terminal EGFP-tagged Fpn where the tyrosine residue position 64 was mutated to histidine using whole-plasmid PCR and ligation (Q5 SDM, NEB) | This study, 37-38 |
| Fpn-GFP (D39A) | C-Terminal EGFP-tagged Fpn where the aspartic acid residue position 39 was mutated to alanine using whole-plasmid PCR and ligation (Q5 SDM, NEB) | This study, 41 |
| Fpn-GFP (D181V) | C-Terminal EGFP-tagged Fpn where the aspartic acid residue position 181 was mutated to valine using whole-plasmid PCR and ligation (Q5 SDM, NEB) | This study, 41, 73 |

**Table S2.**

| Target Gene | Application | Primer Sequence (5' to 3') | Product Size (bp) | Reference |
|---|---|---|---|---|
| Human *Fpn* mRNA | RT-qPCR | F: CGTCATTGCTGCTAGAATCG<br>R: AGACTGAAATCAATACGAGC | 203 | 66 |
| Human *HPRT* mRNA | RT-qPCR | F: CCCTGGCGTCGTGATTAGTG<br>R: GAGCACACAGAGGGCTACAA | 190 | This study |
| *Aph* 16S rRNA | RT-qPCR | F: GGTGAGTAATGCATAGGAATC<br>R: GCTCATCTAATAGCGATAAATC | 108 | 65 |
| Human *HAMP* | RT-qPCR | F: TCCCACAACAGACGGGACAA<br>R: AGCAGCCGCAGCAGAAAAT | 138 | 67 |
| SDM of Fpn-GFP (WT) for **Y64H** Mutation | Cloning (SDM) | F: GACAGCAGTCCATGGGCTGGTGG<br>R: AAAAGGAGGCTGTTTCCATAG | | This study, 37-38 |
| SDM of Fpn-GFP (WT) for **D39A** Mutation | Cloning (SDM) | F: TACTTGGGGAGCGCGGATGTGGC<br>R: GAGAGAGAATGACCAAGGTAG | | This study, 41 |
| SDM of Fpn-GFP (WT) for **D181V** Mutation | Cloning (SDM) | F: ACGAAGGATTGTGCAGTTAACCAAC<br>R: ATTGTGGCATTCATATTTG | | This study, 41, 73 |

Underlined: Sequence for desired mutations
